# The effects of time-varying temperature on delays in genetic networks

**DOI:** 10.1101/019687

**Authors:** Marcella M. Gomez, Richard M. Murray, Matthew R. Bennett

## Abstract

Delays in gene networks result from the sequential nature of protein assembly. However, it is unclear how models of gene networks that use delays should be modified when considering time-dependent changes in temperature. This is important, as delay is often used in models of genetic oscillators that can be entrained by periodic fluctuations in temperature. Here, we analytically derive the time dependence of delay distributions in response to time-varying temperature changes. We find that the resulting time-varying delay is nonlinearly dependent on parameters of the time-varying temperature such as amplitude and frequency, therefore, applying an Arrhenius scaling may result in erroneous conclusions. We use these results to examine a model of a synthetic gene oscillator with temperature compensation. We show that temperature entrainment follows from the same mechanism that results in temperature compensation. Under a common Arrhenius scaling alone, the frequency of the oscillator is sensitive to changes in the mean temperature but robust to changes in the frequency of a periodically time-varying temperature. When a mechanism for temperature compensation is included in the model, however, we show that the oscillator is entrained by periodically varying temperature even when maintaining insensitivity to the mean temperature.

## 1 Introduction

Biochemical reaction rates, as with all chemical reaction rates, are sensitive to changes in temperature and this effect is captured mathematically by the Arrhenius equation [1, 23]. Temperature dependent rates can also alter the dynamics of gene regulatory networks. Previous studies have examined how gene networks behave at various temperatures, and, in general, their dynamics speed up with increasing temperature [30]. For example, the cell doubling time in root meristems of *Zea mays* decreases 21-fold from 3 — 25° C increase in temperature [9]. In addition, time varying temperatures have been shown to impact gene networks. For instance, circadian oscillators can be entrained by time-varying temperatures that cycle with a period close to 24 hours [4, 21, 29, 40].

In models of gene networks, dynamical delay has been used to model the sequential assembly of messenger RNA and then protein. Nucleic acids must be added one by one to the growing mRNA chain, while amino acids are joined end to end with peptide bonds to create a protein. In each case, the large chain of linear reactions can be compactly modeled either with a discrete delay term, or as a distributed delay term [3, 13]. The incorporation of delay greatly simplifies models of genetic oscillators while simultaneously maintaining qualitative similarities to experimental data [6, 16, 34, 35]. Delay-based models play a central role in understanding the origin of oscillations in genetic networks [25, 26] and other nonlinear systems [13, 17]. For constant temperatures, the delay time or distribution can be scaled with the Arrhenius equation, just as the reaction rates. However, less is known about how time-varying temperatures influence delays in such analyses.

Here, we investigate how time-varying temperatures affect delays in genetic networks. We first derive how time-dependent temperature affects the delay term. To do this, we assume that delays arise from a sequence of first-order reactions that can be modeled as an aggregate delay. Although delay in gene networks is the result of the sequential assembly of first mRNA [5] and then protein [27] (see Fig. 2.1), we lump these delays into one term and represent protein production delay as a reduction of a linear chain of reactions. Each reaction in the sequence is then scaled by a common time-dependent Arrhenius factor. Since changes in temperature influence each biochemical step in the sequence that constitute the delay, the value of the delay time will change. From these assumptions, we derive an expression for the time-dependent distribution of delay times. We analyze changes in phase shift and amplitude of the resulting time-varying delay as a function of parameters of a sinusoidally time-varying rate-coefficient induced by temperature changes. We find a nonlinear relationship and, furthermore, find specific cases for which a delay can remain approximately time-invariant under time-varying conditions.

**Figure 2.1.**
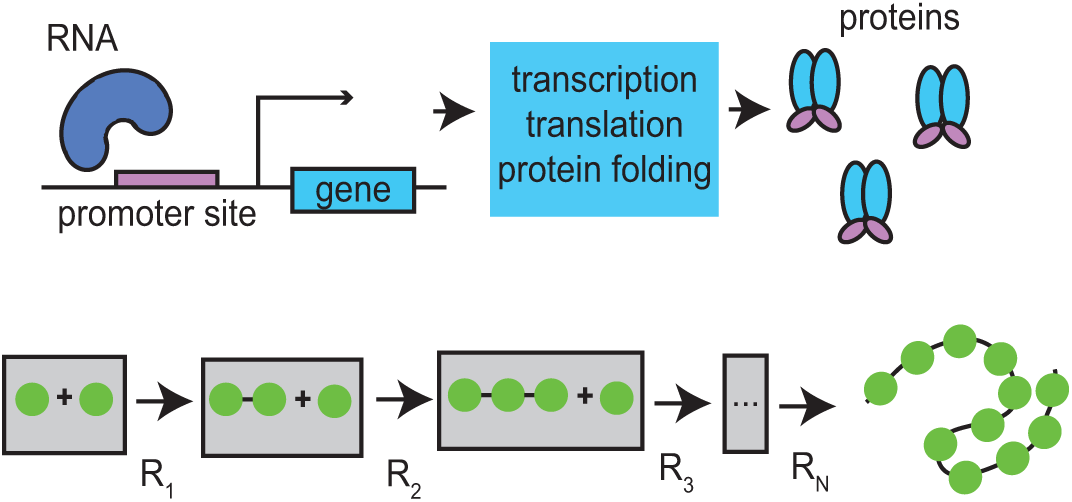
Modeling of delays in protein production. *Transcriptional delays (top) are modeled by a sequence of chemical reactions (bottom) with common reaction rate a*(*t*) *for each reaction R*_1_,..., *R_N_.*

The effects of temperature on oscillators becomes important in the study of circadian clocks and is typically inferred through analysis of system response to single step changes in temperature or a single cycle [21]. We incorporate our findings into a model of a synthetic gene oscillator with temperature compensation presented by Hussain *et al.* [16]. We find that, when the temperature varies sinusoidally in time, the oscillator can be entrained by temperature, but that this entrainment does not occur in the absence of the temperature compensation mechanism. In other words we find that the temperature compensated oscillator is insensitive to changes in the mean temperature but also entrained by periodically varying temperatures, a property explained and observed in circadian oscillators.

## 2 Characterization of time-varying delays

We begin by approximating protein production with a linear sequence of reactions (see Fig. 2.1), the dynamics of which can be modeled by

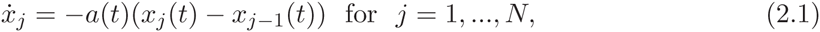

where *x_i_*(*t*) is the concentration of the *i*^th^ species at time *t*, *x*_0_(*t*) is the time varying concentration of the initial complex, *a*(*t*) is the time-varying rate coefficient, and the overdot represents differentiation with respect to time. The effects of time-varying temperatures can be reflected in the time-varying rate coefficients. From this we deduce the effects of time-varying temperatures on the delay distribution, *i.e.* the time it takes to go from the initial complex, *x*_0_, to mature protein, *x_N_*.

To find the distribution function, we first rewrite system (2.1) as

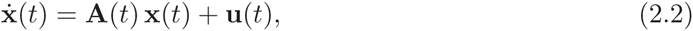

where **x**(*t*) = [*x_N_*,*x_N_* _−1_,...,*x*_1_]*^T^*, **u**(*t*) = [0,..., 0, *a*(*t*)*x*_0_ (*t*)] and **A**(*t*) = *a*(*t*) **J**_−1,_*_N_*. Here

**J**_−1_,*_N_* is the *N*-dimensional Jordan matrix with eigenvalues −1,

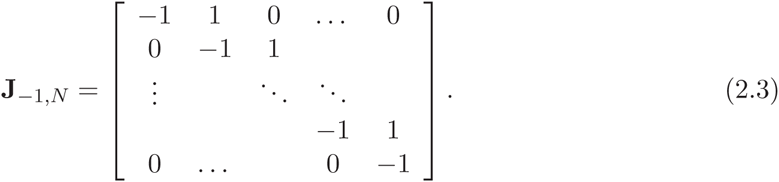

Because **A**(*t*_1_) commutes with **A**(*t*_2_) for all (*t*_1_, *t*_2_), we can write the general solution to equation (2.2) as

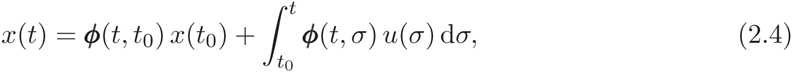

where

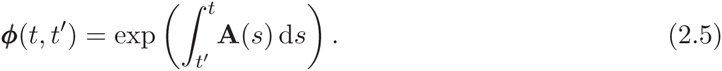

Without loss of generality we set *t*_0_ = 0 and substitute *σ* = *t* − *τ*. If we assume *x_j_*(*t*_0_) = 0 for *j* = 1,..., *N*, the solution reduces to

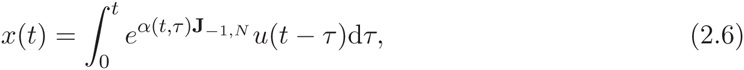

where 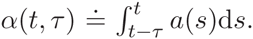. We can now extract the expression relating the input, *x*_0_(*t*), to the measured output, *x_N_*(*t*), with the result

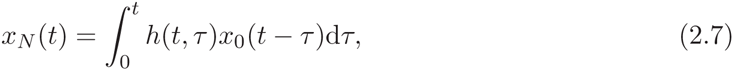

where the function

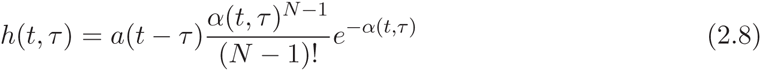

is the impulse response function relating the output to the input of the system, which represents the delay distribution given the constraint 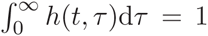, for any time *t*. This constraint is always satisfied by the physics of the problem. The integral can be shown to equal one when *α*(*t* − *τ*) > *ε* for some *ε* > 0, that is, the reaction rate at all times is positive definite. To show this we express the integral as a line integral:

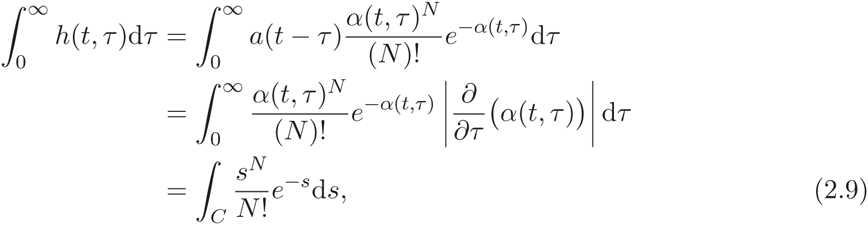

where the curve *C* is the domain of integration that is defined by *α*(*t*, *τ*) for *t* held fixed. Note that the expression in the last line is an integral over the Erlang distribution, which is equal to one when integrated along the curve *C* ≡ *ατ*. For the line integral in equation (2.9) to equal one, *α*(*t*, *τ*) must be an injective function in *τ* meaning d*α*(*t*, *τ*)/d*τ* = *a*(*t* − *τ*) > 0 (which implies *a*(*t*) > 0 for all *t*) with *α*(t, 0) = 0 and lim*τ*_→∞_ *α*(*t*, *τ*) = ∞. By the definition of 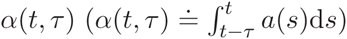, we see that a positive definite *a*(*t*) in turn satisfies the latter conditions.

For the purpose of demonstration we consider a sinusoidally time-varying rate coefficient *a*(*t*) = *δ_p_ a*_0_ sin(*ω t*) + *a*_0_ with 0 < *δ_p_* < 1, assuming the dynamics are induced from an appropriate time-varying temperature. Next, we show conditions under which this is a good approximation for a sinusoidally time-varying temperature. When *a*(*t*) ≡ const. (*i.e. a*(*t*) = *a*_0_), equation (2.8) is the *Erlang* distribution [24]. If *a*(*t*) is not constant, the delay distribution will be a function of time. Figure 2.2(a)-(b) shows the delay distribution *h*(*t*, *τ*) for different values of *N* (holding *N*/*a*_0_ constant), comparing the time-invariant case to the time-varying case. Note that, unlike the time-invariant case (Fig. 2.2(a)), the distribution in the time-varying case (Fig. 2.2(b)) need not be unimodal, especially for small *N*. Figure 2.2(c) shows how the time-varying distribution changes with time for fixed *N* and a time-varying rate coefficient *a*(*t*). Note that the distribution becomes unimodal as *N* increases.

**Figure 2.2.**
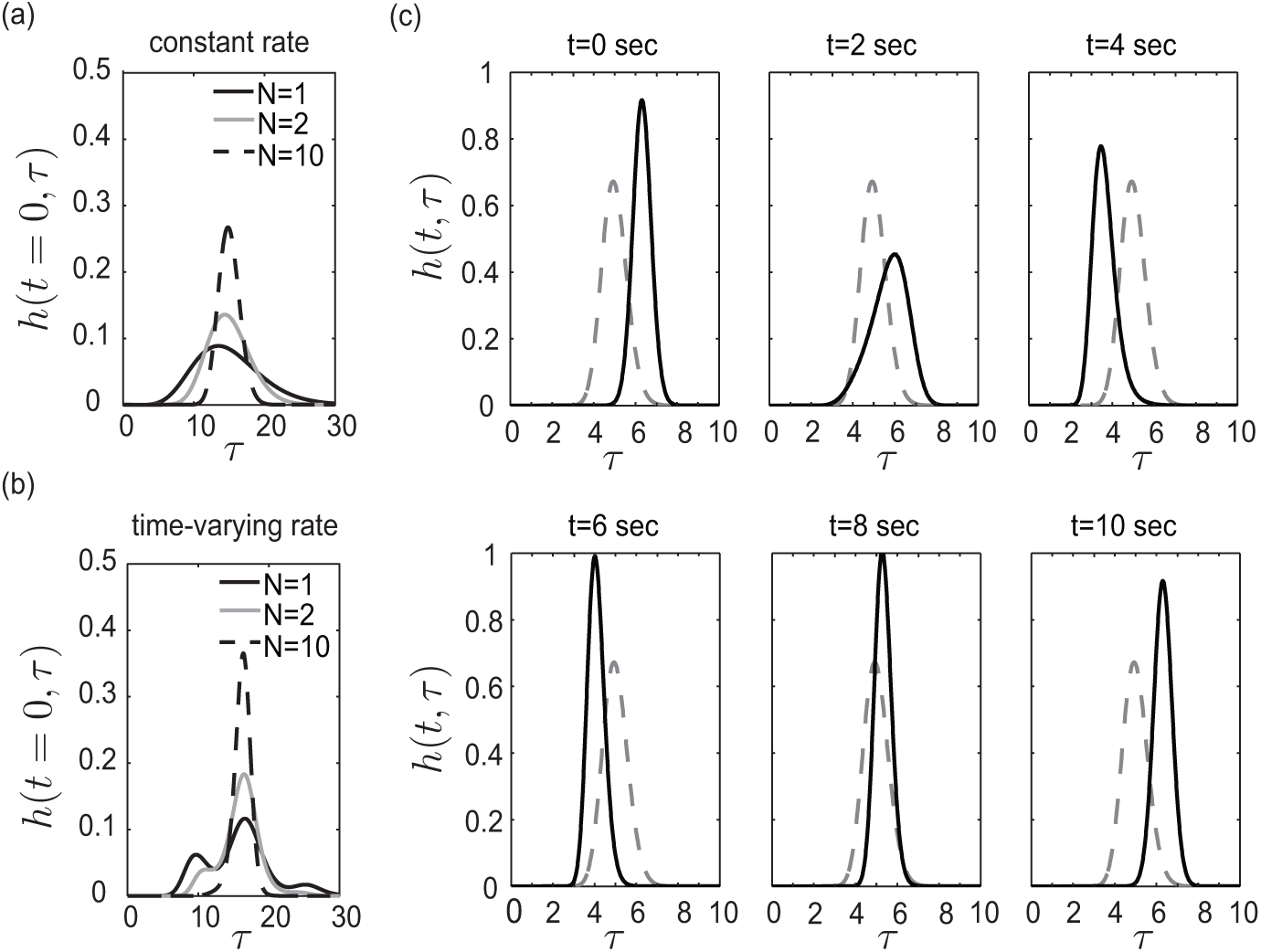
Delay distribution for different values of *N* with *E* = 15 for a constant and time-varying rate *a.* *(a) Delay distribution for different values of N with E* = 15 *for a constant rate coefficient a. (b) Delay distribution for different values of N with E* = 15 *for a time-varying rate coefficient a(t)* .5 *a*_0_ sin(ω *t*) + *a*_0_ *at time t* = 0. (*c*) *Distribution as a function of time with a*(*t*) = *a*_0_ *δ_p_* sin(*ω t*) + *a*_0_*, N* = 100, (*N* + 1)/*a*_0_ = 15, δ*_p_* = .5, *and ω* = 2*π/*20. *The dashed line indicates the nominal time-invariant distribution with a*(*t*) = *a*_0_.

Next, consider the limit as the number of reactions within the sequence tends to infinity. In the time-invariant case, one would consider the limit as *N* → ∞ such that *N/a* = *E* remains constant. Taking this limit reduces the distribution function to the Dirac delta function *δ*(*τ* − *E*), which agrees with results shown by Bel *et al.* [3]. To investigate the time-varying case, we assume that *a*(*t*) = *a*_0_ *f* (*t*), where *a*_0_ > 0 and *f*(*t*) is a positive definite, bounded function of time, which agrees with the sinusoidally varying function *a*(*t*) above. In this case, if we take the limit *N* → ∞ with the constraint *N*/*a*_0_ = *E*, the ratio *N*/*α*(*t*, *τ*) remains finite for finite *τ*.

In summary, we find that there exists a unique delay *τ*_eff_ such that

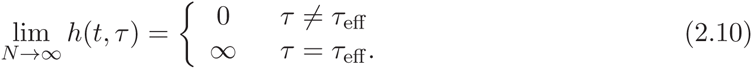

The derivation of these results can be found in Appendix A. Therefore, with the integral over the function equal to one, in the limit as *N* → ∞ such that *N*/*a*_0_ = *E*_0_ the distribution is approximated by a delta function centered at *τ*_eff_ (*i.e.* lim*_N_*_→∞_ *h*(*t*, *τ*) ~ *δ*(*t* — *τ*_eff_(*t*))), which is necessarily a function of *a*(*t*), and therefore time-varying. From the derivation we find that this unique delay *τ*_eff_(*t*) can be found by solving

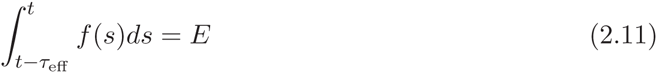

for *τ*_eff_. Note that *τ*_eff_(*t*) can be computed with only the expected delay *E* = *N*/*a*_0_ and the time-varying function *f*(*t*). Also, since *f*(*s*) is positive definite, we can guarantee a single solution *τ*_eff_ for every time *t*. The effective delay *τ*_eff_ is computed such that the integral remains constant at *E*. Note that the area under the curve is zero for *τ*_eff_ = 0 and monotonically increases with increasing *τ*_eff_.

We now apply the method to investigate delays under periodically time-varying temperatures. For a time-varying temperature

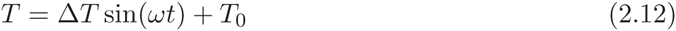

we can rewrite the Arrhenius equation *A*(*T*) = *A*_0_ *e*^−^*^θ/T^* in the form

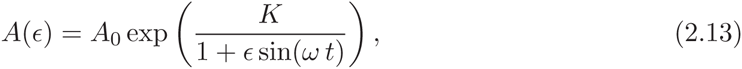

where 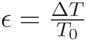 is non-negative and *K* = −*θ/T*_0_. We assume *ε* ≪ 1 (i.e. Δ*T* ≪ *T*_0_) and in the Taylor expansion we have

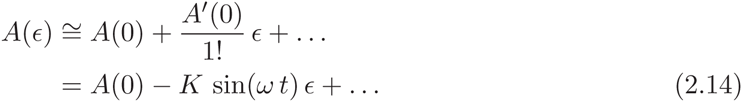

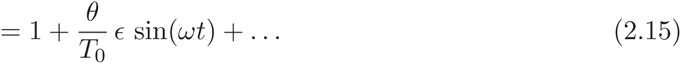

Here, *A*_0_ is chosen such that *A*(0) = 1, *i.e. A*_0_ = exp(*θ*/*T*_0_). Therefore for small *ε* we can approximate the time-varying Arrhenius equation by

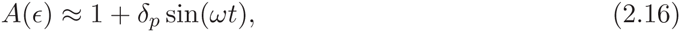

where 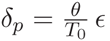. The time-varying rate coefficient for the reaction rates implicit in the delay are given by *a*(*t*) = *a*_0_ · *A*(*t*).

For a sinusoidally varying rate coefficient *a*(*t*) = *a*_0_ *δ_p_* sin(*ωt*) + *a*_0_, using equation (2.11), the effective delay reduces to solving

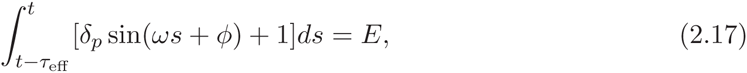

where *E* is the expected delay. In the limit analyzed, changes in the delay are determined only from the expected delay without perturbation and the perturbation on the reaction rates. Also note that in the extreme limits of the frequency, we have

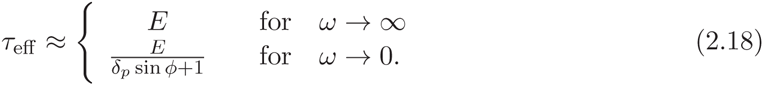

Taking the integral in equation (2.17), the solution can be shown to solve a function of the form

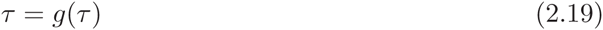

where

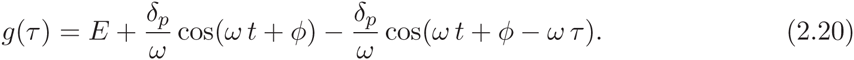

In this case the effective delay at each time *t* must be solved numerically. Figure 2.3 shows the delay as a function of time for a periodically varying temperature. The solution is found numerically for a discretized range of time. In Fig. 2.3 we consider the expected delay *E* = 13.5 min., which is what is chosen for *τ_x_.*

**Figure 2.3.**
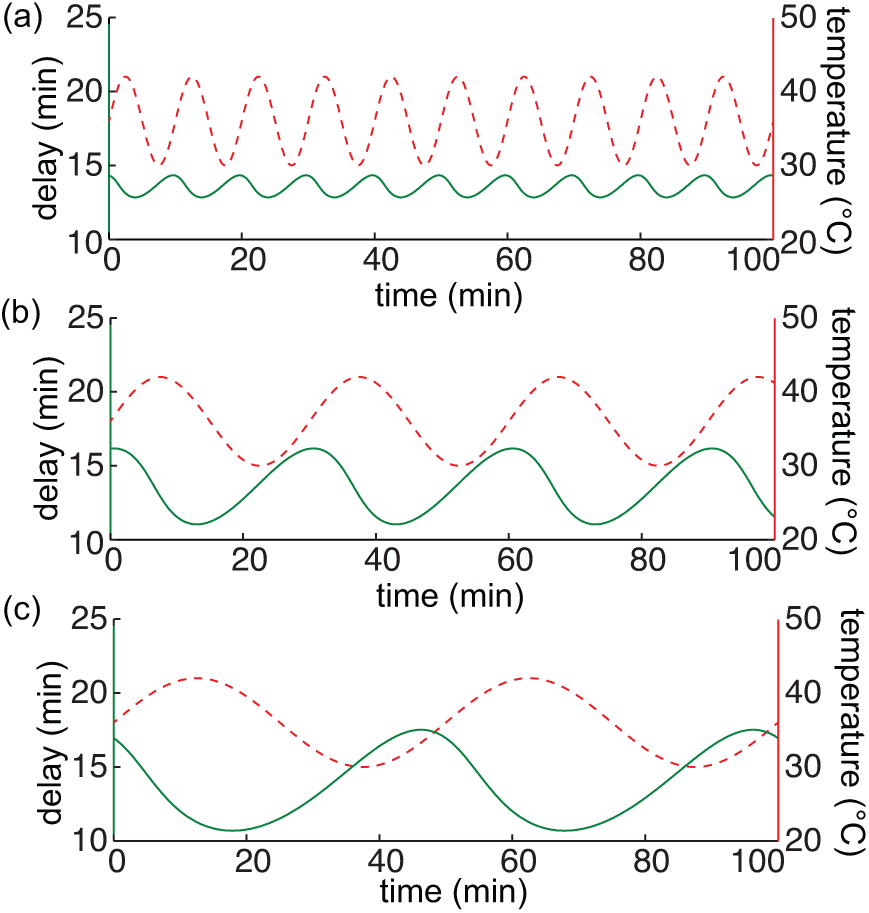
Amplitude of the delay changes with the period of the time-varying temperature. *Time-varying delays (green solid lines) corresponding to various time-varying temperature T*(*t*) = Δ *T* sin(*ωt*) + *T*_0_ *(red dashed lines). Parameter values here are θ* = 4500*K, E* = 13.5 *min.,* Δ*T* = 6° *C, T*_0_ = 36° *C, and δ_p_* = .28. *(a)* 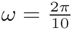. *(b)* 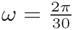. *(c)* 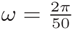.

We now consider the effects of changing parameters *δ_p_* and *ω* on the time-varying delay. Figure 2.4 shows analysis of the calculated time-varying delay as a function of various parameters. Figure 2.4(a) shows an example of the time-varying function *τ*_eff_ (*t*). In Figure 2.4(b)-(d), we look at how the mean and amplitude of the time-varying function *τ*_eff_ changes as we change the expected delay *E*, the relative perturbation *δ_p_*, and the frequency *ω* of the time-varying rate function *a*(*t*). Most of the results are in line with intuition. For example the amplitude of *τ*_eff_(*t*) increases with an increase in the relative perturbation *δ_p_* and decreases as the frequency *ω* of *a*(*t*) increases. If the environmental conditions change too quickly, the system does not effectively respond. An unexpected result is the non-monotonic behavior of the function *τ*_eff_(*t*) as the mean delay changes. Figure 2.4(b), implies that *τ*_eff_ remains constant when the mean delay is exactly equal to the period of *a*(*t*). With this observation we note that if *ω* = 2*πn*/*T* for any positive integer n in equation (2.19),(2.20), then we always have the solution *τ*_eff_(*t*) = *T*. This suggests that delays are minimally affected by sinusoidally time-varying reaction rates when the mean delay is an integer multiple of the period. We see the the result of this in Figure 2.4(c) as well.

**Figure 2.4.**
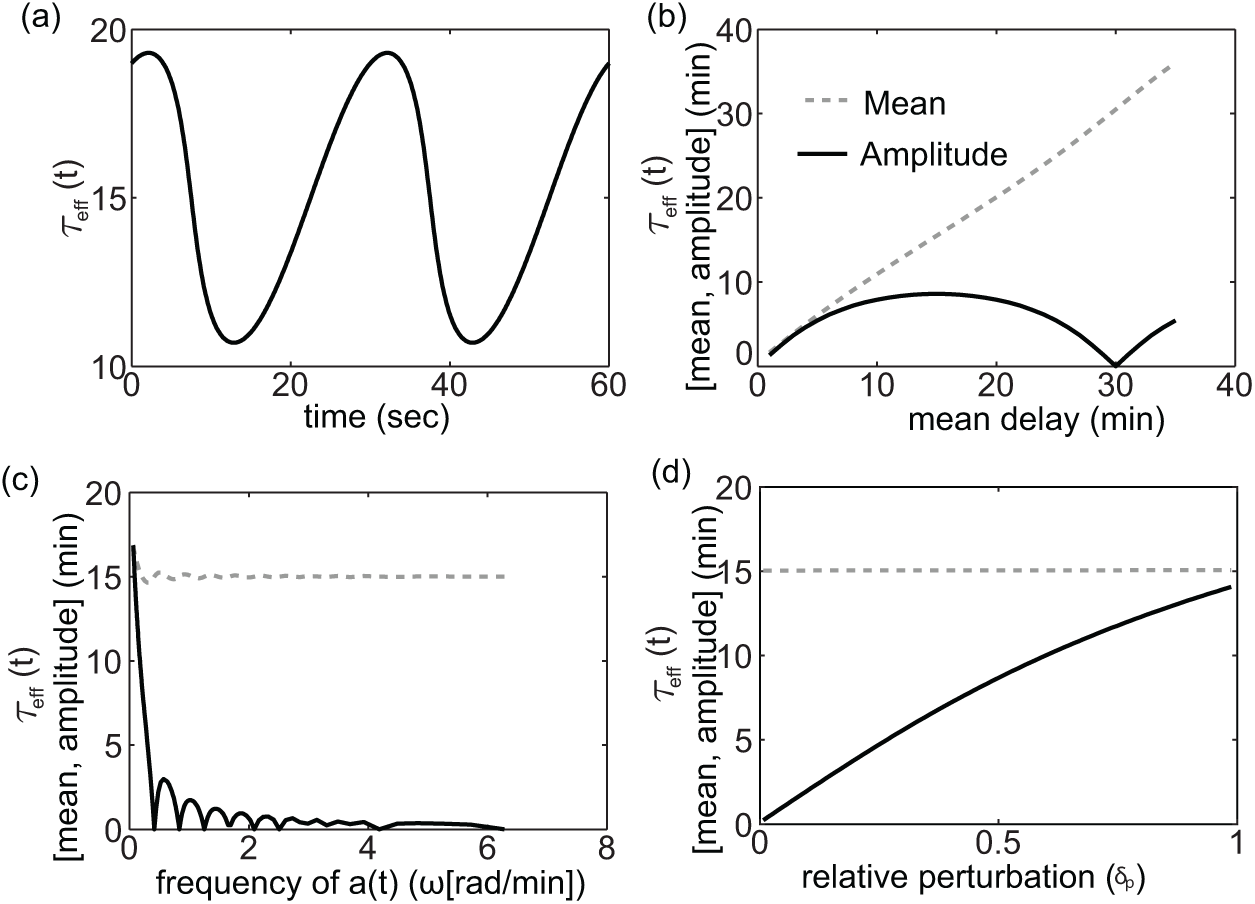
Mean and peak-to-peak amplitude of *τ*_eff_ as a function of *δ_p_*, *E*, and *ω*. *(a) τ*_eff_ *as a function of time with ω = π*/15, *φ* = 0, *E* = 15, *and δ_p_* = .5. *(b) τ*_eff_ *as a function of the mean delay with ω = π*/15, *φ* = 0, *and δ_p_* = .5. *(c) τ*_eff_ *as a function of the frequency of a*(*t*) *with φ* = 0, *E* = 15, *and δ_p_* = .5. (*d*) *τ*_eff_ *as a function of the relative perturbation δ_p_ with ω = π/E, φ* = 0, *and E* = 15.

It is apparent from Figure 2.4, that the dependence of *τ*_eff_ on the sinusoidally time-varying temperature can be nonlinear. In this respect, we analyze the phase shift between the sinusoidally time-varying reaction rate *a*(*t*) and the resulting time-varying delay *τ*_eff_(*t*). In Figure 2.5(a)-(c), we look at how the phase shift changes as we change the mean delay *E*, the relative perturbation *δ_p_*, and the period of the time-varying rate function *a*(*t*). Before calculating phase shift we account for the fact that the reaction rate and delay functions are initially 180° out of phase because the delay decreases when the reaction rate *a*(*t*) increases. Also, since *τ*_eff_(*t*) is not a perfect sinusoid, we calculate phase shift based on the distance between peaks. In general we find an increase in phase shift with an increase in mean delay *E*, the relative perturbation *δ_p_*, and the period of the time-varying rate function *a*(*t*). However, Figure 2.5(b)-(c) show an existence of discontinuities in the phase shift. There is a 180° phase jump when the mean delay equals the period. Recall, that the amplitude of *τ*_eff_(*t*) becomes zero when delay is an integer multiple of the period. As the response *τ*_eff_(*t*) crosses this critical point there is a 180° phase shift as the amplitude of the response becomes non-zero again. Furthermore, the frequency of discontinuities increase on the log scale as a function of frequency and mean delay. We show trends in a limited range of frequencies and mean delay in Figure 2.5(b)-(c) to help highlight where the nonlinearities come from and demonstrate the non-trivial behavior.

**Figure 2.5.**
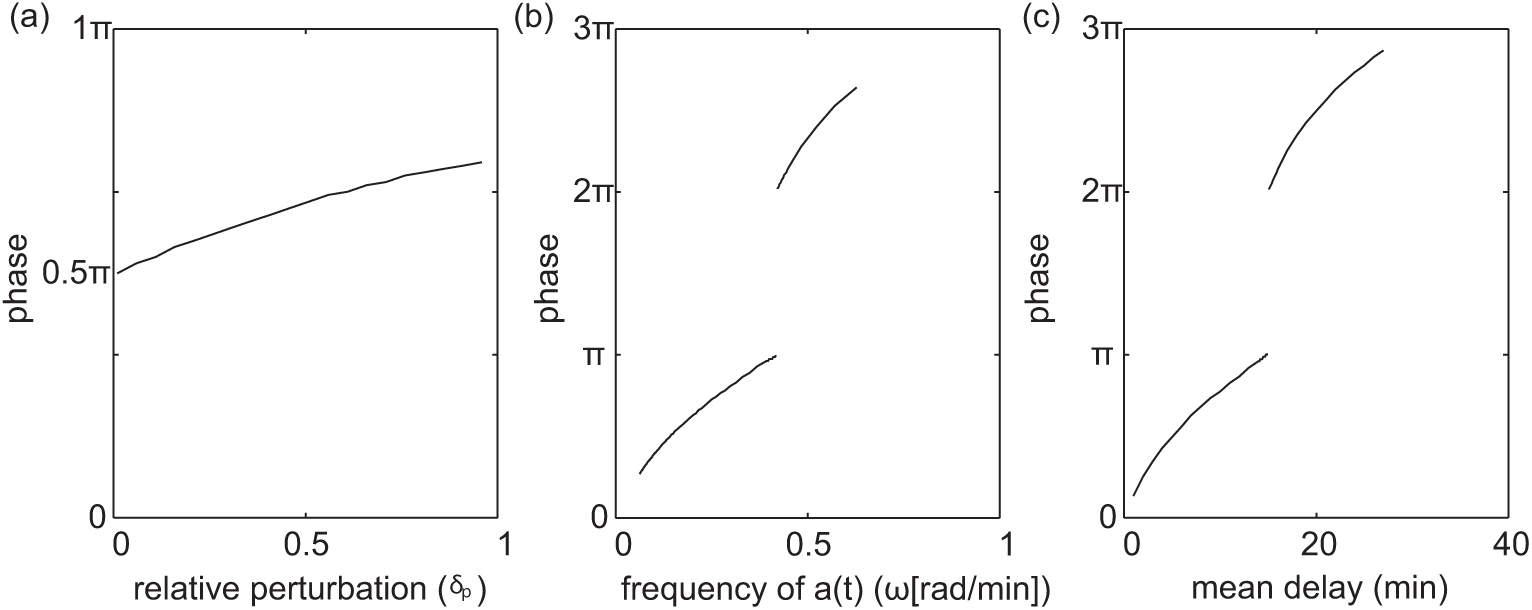
Phase shift between *a*(*t*) and *τ*_eff_(*t*) as a function of *δ_p_*, *E*, and *ω*. *(a) phase shift as a function of δ_p_ with E* = 15 *and ω* = 2*π*/30. *(b) phase shift as a function of ω with E* = 15, and *δ_p_* = .5. *(c) phase shift as a function of E with ω* = 2*π*/50, and *δ_p_* = .5.

## 3 Temperature Entrainment of a Dual-Feedback Oscillator

We now consider the entrainment properties of a temperature compensated dual-feedback oscillator presented by Hussain *et al.* [16]. The oscillator, as depicted in Fig. 3.1, can be modeled as [16]:

**Figure 3.1.**
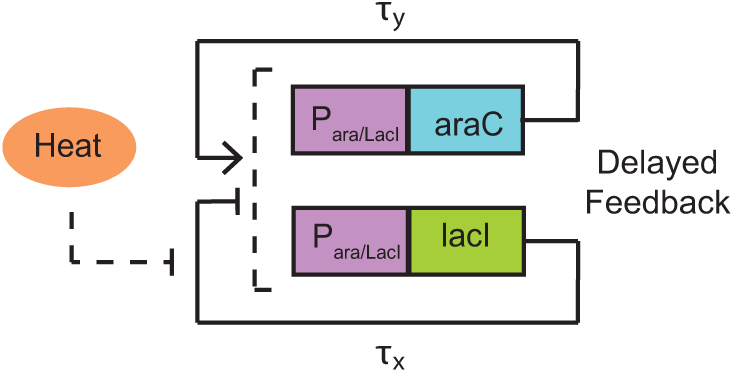
Schematic of the temperature compensating oscillator [16].

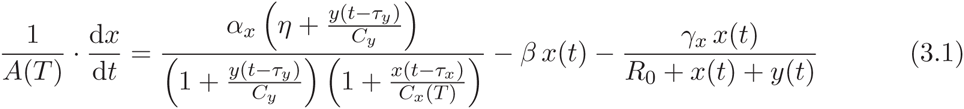

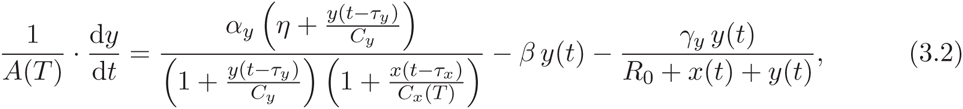

where *x* and *y* are the concentrations of the repressor (LacI) the activator (AraC); *α_x_* and *α_y_* are the maximal transcription initiation rates for *x* and *y*, respectively; *C_x_* and *C_y_* are the binding affinities of LacI and AraC to the promoter, respectively; *β* is the dilution rate due to cellular growth; *η* is a measure of the strength of the positive feedback loop; *R*_0_, *γ_x_*, and *γ_y_* are Michaelis-Menten constants for enzymatic decay of the proteins; *τ_x_* an *τ_y_* are temperature dependent delay times for the production of LacI and AraC, respectively; and *A*(*T*) is the common Arrhenius scaling of all reaction rates. Additionally, the Arrhenius scaling term has the form *A*(*T*) = *A*_0_ *e*^−^*^θ^*^/^*^T^*, where *θ* is the temperature scale. Note that increasing temperature increases the scaling coefficient *A*(*T*) and hence speeds up the dynamics of the system. In Hussain *et al.* [16], the authors scale the delay by the Arrhenius constant when predicting dynamics at varying temperatures (the temperatures are held constant for each assay). In this case, we consider predictions under time-varying temperatures defined by equation (2.12). We use the method derived in Section 2 to determine the resulting time-varying delay. The binding affinity of LacI, *C_x_*(*T*) is a also a function of the temperature

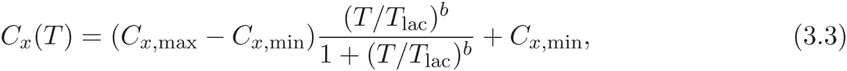

and provides the mechanism for temperature compensation in the oscillator [16]. *C_x_*_,min_ and *C_x_*_,max_ are the minimum and maximum biding affinities of LacI to its promoter. *T*_lac_ is the temperature at which *C_x_*(*T*) is half-maximal and *b* is a Hill coefficient.

In Hussain *et al.* [16], the period of the oscillator is shown to remain largely unaffected by changes in constant temperature due to a temperature sensitive LacI mutant, which is modeled by a temperature dependent binding affinity *C_x_*(*T*). In order to compare entrainment properties to a system without such a temperature dependent mechanism, we consider a similar model where the binding affinity *C_x_*(*T*) = *C_x_*(*T*_0_) remains constant. In this case temperature affects are introduced solely through an Arrhenius scaling and implicitly through the time-varying delay. Given equation (2.16), we drive the system with a time-varying temperature described by equation (2.12) with *θ* = 4500*K* and Δ*T* = 2°*C*. Details of the simulations are found in Appendix B. In Fig. 3.2(a) we fix the frequency at 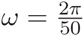 rad/min and vary the mean temperature *T*_0_ in order to verify the temperature compensating property achieved through the temperature dependent LacI mutant. Without the temperature compensating mechanism the frequency of oscillations changes linearly with the mean temperature but remains constant with the LacI mutant. In Fig. 3.2(b) we fix the mean temperature at *T*_0_ = 36°*C* and vary the frequency *ω* to study frequency entrainment for the same system with and without the temperature compensating mechanism. It is clear that the system entrains only under the influence of the temperature sensitive promoter. A common Arrhenius scaling alone does not allow for frequency or phase entrainment. The same mechanism that provides temperature compensation (insensitivity to changes in mean temperature) also makes the system sensitive to temperature dynamics, achieving entrainment. This is in agreement with circadian clocks as well [4].

**Figure 3.2.**
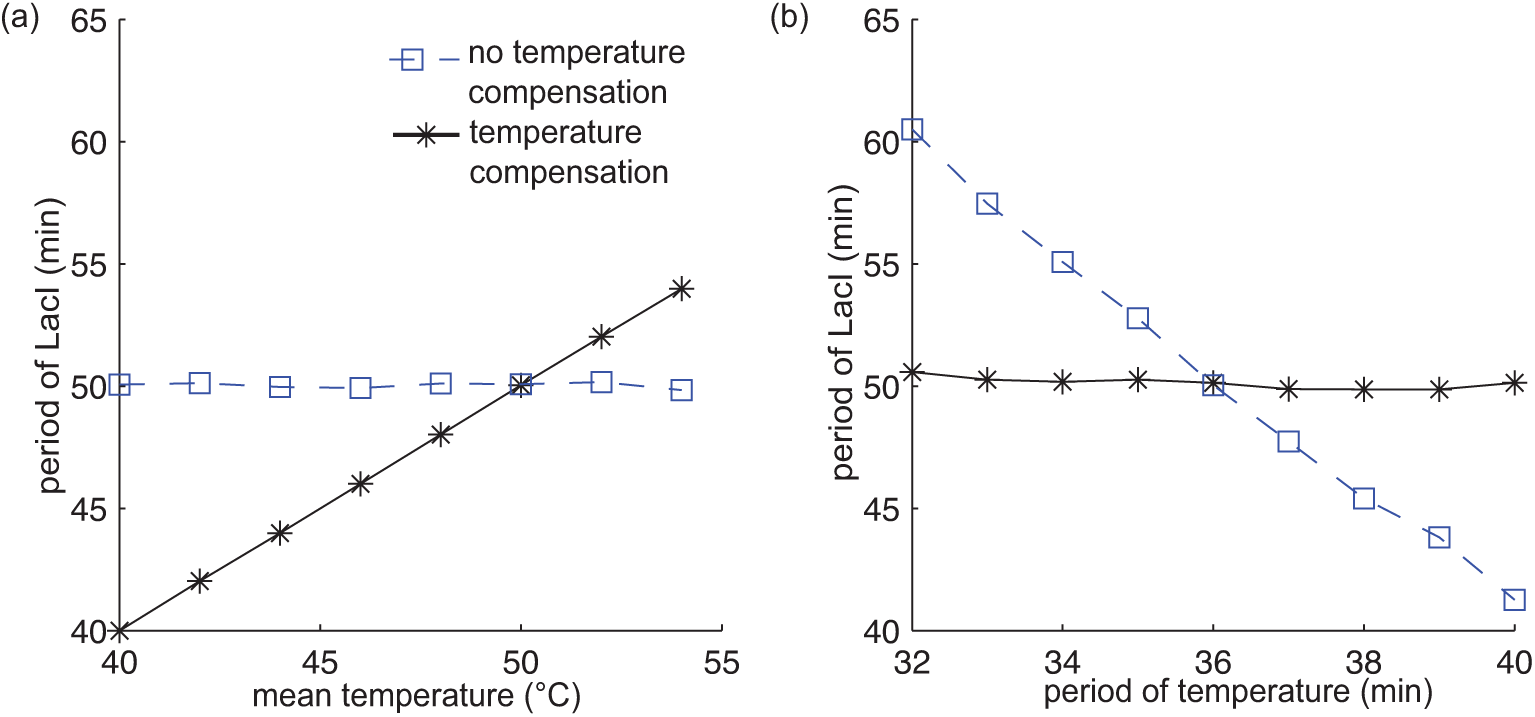
Entrainment of the synthetic gene oscillator with and without a temperature sensitive promoter. *(a) Period of the circuit with and without temperature compensation for ω* = 2*π*/50min^−1^, Δ*T* = 2°*C, and different mean temperatures T*_0_ *in Eq. (2.12). (b) Frequency entrainment of the circuit with and without temperature compensation for* Δ*T* = 2° *C and T*_0_ = 36° *C.*

## 4 Conclusion

It was found that periodic temperature fluctuations induce periodically time-varying delays. The effects of a time-varying temperature on delays within a genetic network can be highly nonlinear and so the delay cannot be simply scaled by an Arrhenius coefficient in this case. With this, we investigated properties of a delay-based model of a temperature sensitive oscillator. This oscillator has been shown to exhibit temperature compensation, that is, the frequency of oscillation is insensitive to temperature variations. This was shown by analyzing the dynamics at different constant temperatures. Using the method derived, we were able to simulate the system under a periodically time-varying temperature. Simulations showed improved temperature compensating properties under the dynamically varying temperatures, over constant temperatures. Simulations also predicted reliable temperature entrainment. The frequency of protein expression coincided with that of the time-varying temperature.

We focused on properties important in circadian oscillators, namely, temperature compensation and temperature entrainment. Ideally, a circadian oscillator should demonstrate properties of entrainment with insensitivity to changes in mean temperatures [8]. Here we highlight a case where the entrainment is a byproduct of the same mechanism which makes the system insensitive to changes in mean temperature. This is in agreement with Bodenstein *et al.* [4], where temperature entrainment was shown to naturally follow from circadian clock models tuned for temperature compensation through the Arrhenius coefficients. In the oscillator of Hussain *et al.* [16], there is an inherent tradeoff between robustness to unwanted temperature fluctuations about a mean and robustness to changes in mean temperatures, with the latter admitting temperature entrainment. Here, an understanding of the effects of temperature on delays eased the analysis of a delay-based model of a circuit with circadian clock-like properties.

Future work includes investigation of circadian oscillators, which have an intricate relationship with temperature. For instance, circadian oscillators exhibit temperature compensation [2, 31], *i.e.* their periods do not vary with changes in the average temperature. Theorists have investigated methods of temperature compensation in models of circadian oscillators, often minimizing the effects of Arrhenius-scaled rate constants [8, 12, 14, 15, 33, 36]. Periodic changes in temperature have also been implicated in the entrainment of circadian oscillators to the day/night cycle [21, 29, 32, 37, 40]. However, entrainment of circadian oscillators is most commonly associated with periodic changes in light, and mathematical models have been developed explaining this phenomenon [7, 10, 11, 18, 19, 20, 22, 28, 38, 39]. Less is understood about the role of temperature.

## Appendix A Time-varying distribution limit

Here, we provide details on taking the limit *N* → ∞ with the constraint *N*/*a*_0_ = *E* on the time-varying distribution

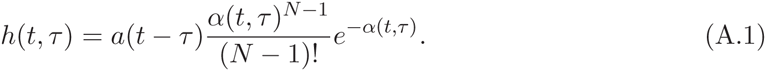

We will show that in the limit the distribution, while maintaining an integral equal to one (as shown in the main text), becomes zero everywhere and infinity at a single point. In summary, we find that there exists a unique delay *τ*_eff_ such that

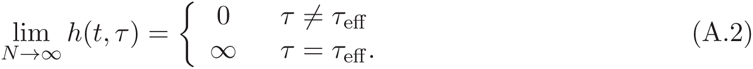

Applying Stirling’s formula for large *N*, namely

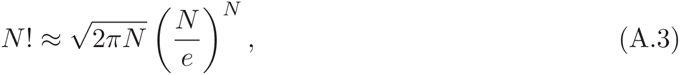

to the distribution (A.1) gives

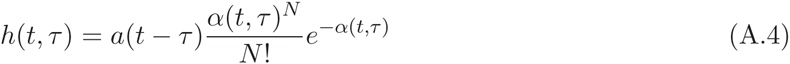

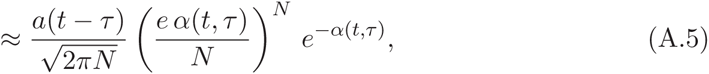

which asymptotically converges to equation (A.1) in the limit as *N* → ∞. Rearranging terms in equation (A.5) and making use of the substitution *a*(*t*) = *a*_0_ *f*(*t*) gives

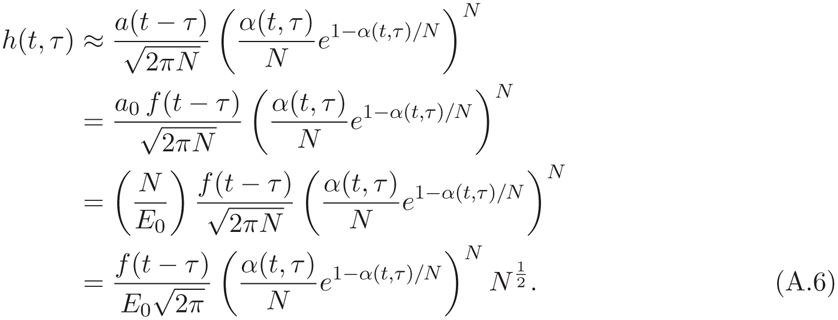

We define

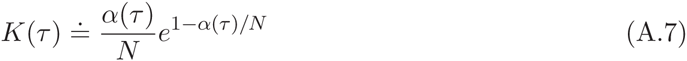

and investigate the limit of equation (A.6) for different ranges of *K* by looking at the term

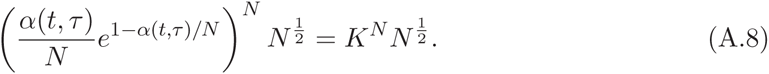

Note that *K* remains constant with changing *N* under the constraint *N*/*a*_0_ = *E*. For ease of analysis we ignore the coefficient 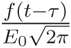 in equation (A.6), which also does not change in the limit. We will show that in the limit N → ∞, equation (A.8) is zero everywhere and infinity at a singular point for any time *t*.

Applying l’H*τ*ptial’s rule for *K* < 1

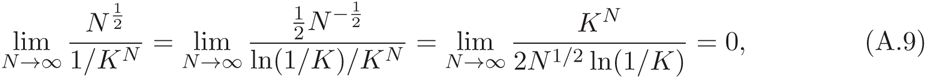

and for *K* ≥ 1

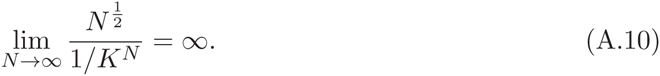

It remains to show that *K* ≤ 1 for all *τ*. We would like to determine when *K* reaches its maximum value. As a necessary condition for an extremum we must have

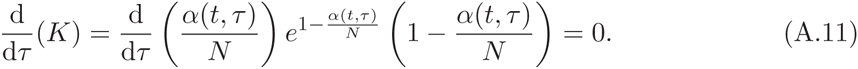

Since the first two terms are always strictly positive, we find that an extremum occurs at *τ*_eff_, where

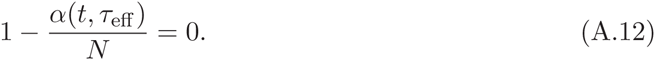

Plugging equation (A.12) back into equation (A.7),

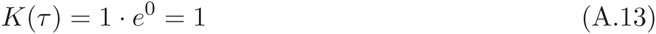

we find *K* = 1 at the extremum. It can be easily shown that

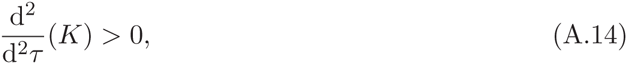

therefore, the extremum is a maximum, hence, *K* ≤ 1 for all *τ*. We see that in the limit as *N* → ∞, *h*(*t*, *τ*) is zero everywhere for all *τ* except for at *τ*_eff_, where *K*(*τ*) = 1 and *h*(*t*, *τ*) = ∞. Furthermore, since *α*(*t*, *τ*) is an injective function in *τ*, equation (A.12) has a single solution, hence, *τ*_eff_ provides a global maximum at a given time *t*, which is the single non-zero solution.

## Appendix B Simulations of the dual-feedback oscillator

The oscillator modeled by

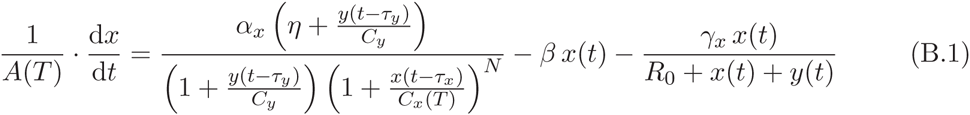

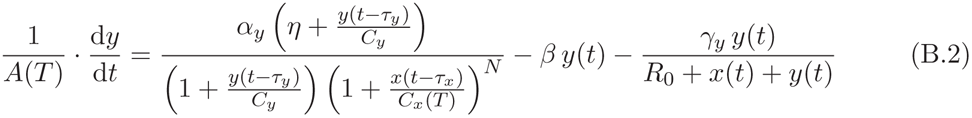

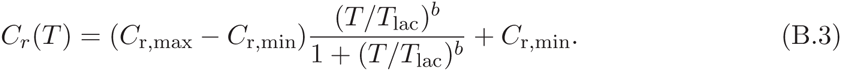

**Table B.1.**
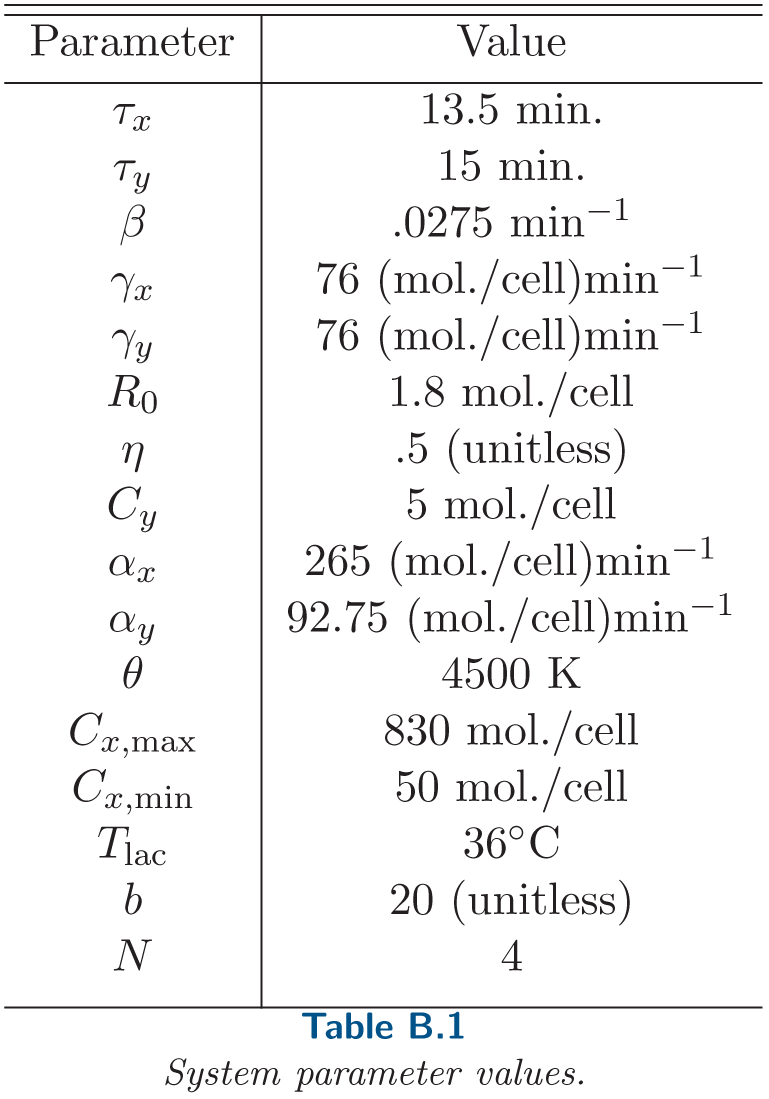
System parameter values.

This system is modeled using dde23 in Matlab with discretized delays. Using dde23, one can specify the delayed states to be used in the delay differential equation. Using the method derived in Section 2, we can determine the range of delays for which we need state information. The range of delays is discretized into bins of width .05min so that state information is saved for N ∈ ℤ different delays, where *N* = (*τ*max − *τ*min)/.05. The delay used in the simulation is the mid-point of each bin. At each iteration, we calculate what the time-varying delay is at that time using again the method in Section 2 and find the appropriate bin. The corresponding delayed state is then fed into the delay differential equation, simulating the time-evolution of the model above with a time-varying delay.

## Acknowledgments

The authors thank Jae Kyoung Kim for insightful discussions

